# c-Fos-MMP-9 pathway in central amygdala mediates approach motivation but not reward consumption

**DOI:** 10.1101/2020.04.17.044792

**Authors:** T. Lebitko, K. Nowicka, J. Dzik, D. Kanigowski, J. Jędrzejewska-Szmek, M. Chaturvedi, T. Jaworski, T. Nikolaev, T. Gorkiewicz, K. Meyza, J. Urban-Ciecko, L. Kaczmarek, E. Knapska

## Abstract

Although impaired motivational and consummatory aspects of reward behavior are core symptoms of several psychiatric disorders, the underlying neural and molecular mechanisms remain poorly understood. c-Fos, as a component of AP-1 transcription factor, regulates the expression of matrix metalloproteinase 9 (MMP-9), an enzyme involved in synaptic remodeling and plasticity. Both proteins are expressed in the central amygdala (CeA) that orchestrates appetitive and aversive responses. We have examined the role of c-Fos and MMP-9 in CeA in reward and punishment processing. We have manipulated c-Fos and MMP-9 levels in vivo using: RNAi-based approach to block c-Fos expression, inhibitor-releasing nanoparticles to block MMP-9 activity, and lentiviral vector to increase MMP-9 expression. To assess motivation, consumption and learning reinforced by either a sweet or bitter-tasting solution, we employed automated behavioral tests in the IntelliCage cystem. We have used transgenic mouse lines to identify CeA cells expressing c-Fos and a specific fluorescently labelled substrate peptide to measure MMP-9 activity. Excitability of neurons was measured with patch-clamp. Blocking c-Fos expression has been found to disrupt both reward processing components, motivational and consummatory, while inhibition of MMP-9 activity has impaired only motivational aspects of the behavior. On the other hand, punishment processing has not been affected by those manipulations. Furthermore, we have observed that reward training induces c-Fos expression in both somatostatin (SST)+, and SST-neurons, while MMP-9 activity is increased in SST- subpopulation only. We have further linked SST+ population to consummatory reactions by showing that reward consumption increases excitability of the SST+ neurons. These findings reveal molecular mechanisms of motivational anhedonia, linking it to c-Fos-MMP-9 pathway and CeA SST- neurons, and consummatory anhedonia, linking it to c-Fos and CeA SST+ neurons.

## INTRODUCTION

Anhedonia, a core symptom of Major Depressive Disorder [1], has long been defined as reduced ability to experience pleasure. This long-standing definition has recently been re-conceptualized to include different deficits in reward processing, i.e., impairments in the ability to pursue or to experience pleasure [2,3]. Both components of reward processing, seeking of and experiencing pleasure, nowadays most commonly referred to as ‘wanting’ and ‘liking’ [4], are important factors for goal-directed reward-based learning [5]. While the seminal works of Berridge [6] pointed out the existence of different neural substrates of wanting and liking in the brain, our understanding of the underlying neural circuitry and molecular underpinnings is still limited.

One of the brain structures implicated in reward processing is the central nucleus of amygdala (CeA) that orchestrates a wide range of behavioral responses. The CeA is functionally heterogeneous and comprises of various, precisely connected, genetically defined cell populations, which control specific behaviors, including appetitively motivated ones [7]. Our previous results showed an increase in c-Fos expression in the CeA after appetitively motivated learning [8]. Furthermore, we have demonstrated that matrix metalloproteinase-9 (MMP-9, extracellularly operating enzyme) in the CeA is crucial for appetitively, but not for aversively motivated discrimination learning [9]. In particular, we have shown that both the constitutive knock-out of MMP-9 and its local inhibition in the CeA impair appetitive, but not aversive discrimination learning [9]. All the experiments were carried out in the IntelliCages. c-Fos, as a component of AP-1 transcription factor, regulates gene expression of MMP-9, an enzyme involved in synaptic remodeling [10]. c-Fos/AP-1driven MMP-9 expression has been demonstrated in the brain *in vivo* during learning [11], as well as in hippocampal neuronal culture *in vitro*, where a specific BDNF-dependent signaling pathway, including c-Fos/AP-1 activating MMP-9 expression, has been revealed [12]. Furthermore, it has also been shown *in vivo* in the brain that MMP-9 cleaves pro-BDNF converting it to its mature form [13]. In aggregate, c-Fos and MMP-9 form a functional and reciprocal pathway in activated neurons, playing a major role in neuronal plasticity [14].

In the present study we aimed to delineate the role of c-Fos and MMP-9 in the CeA in motivational (related to wanting) and consummatory (related to liking) aspects of reward and punishment processing. To distinguish wanting from liking we used long-running tests carried out in a home cage (IntelliCage), allowing measurement of voluntary behaviors and balancing appetitive and aversive conditions [15]. We tested behavioral changes in mice subjected to either local, CeA-limited, blocking of c-Fos expression (by means of RNAi-based approach, [16]), or local inhibition of MMP-9 activity (with nanoparticles releasing TIMP-1, tissue inhibitor of proteinases 1, that endogenously acts as an inhibitor of the enzyme, [9]). Further, to confirm MMP-9 specificity we used mice with a lentiviral vector induced local overexpression of MMP-9 [17].

We have found that blocking either c-Fos or MMP-9 in the CeA affects processing reward but not punishment. Importantly, we have also discovered that blocking c-Fos expression disrupts both reward wanting and reward liking, while blocking MMP-9 activity impairs wanting, leaving liking intact. Next, we have characterized the CeA neurons activated by the reward-motivated training. We have found that such training induces c-Fos expression in both somatostatin-positive (SST+) and somatostatin-negative (SST-) CeA neurons, while MMP-9 activity is increased in the SST-subpopulation only. Furthermore, we have linked the SST+ population with liking by showing that reward consumption results in increased excitability of the SST+ neurons in the CeA. In aggregate, these findings reveal molecular and neural mechanisms of consummatory anhedonia, linking it to c-Fos and the SST+ neurons, and motivational anhedonia, linking it to c-Fos-MMP-9 pathway and the SST- neurons in the CeA.

## MATERIALS AND METHODS

### Animals

All procedures were carried out in accordance with the European Union and Polish guidelines for the care and use of laboratory animals, and approved by the Local Ethical Committee. 7-8-week-old C57BL/6 female mice were obtained from a local commercial supplier, while strains of transgenic SST-Cre (stock #013044) and Ai14 (stock #007908) mice came from Jackson Labs. Experiments were performed in 1-2 months old female offspring of SST-Cre mice crossed to Ai14. Mice were housed in a 12/12 h dark/light cycle, partially reversed (the dark phase shifted from 20:00–8:00 to 13:00–01:00 or 12:00–24:00 dependent on daylight savings), with food available ad libitum.

### Lentiviral Vectors construction and preparation

Vectors bearing shRNA coding sequences sh_luc and sh_cfos, and SYN-MMP9-GFP (Syn-aMMP9 T2A GFP) and SYN-GFP vectors were produced as previously described [16,17]. All vectors were prepared in the Laboratory of Animal Models in the Nencki Institute.

### Formulation of poly(DL-lactide-co-glycolide) (PLGA) nanoparticles

The PLGA nanoparticles (NPs) containing TIMP-1 (the endogenous MMP-9 inhibitor) or inactive protein (bovine serum albumin, BSA) were prepared as previously described [9,11]. For fluorescent visualization, the proteins were tagged with FITC (Sigma, F7250) before encapsulation.

### Stereotactic injections

Surgery room and surgical instruments were sterilized. Mice received subcutaneous butorphanol injection (0.2 mg/100 g), were placed on a heating pad and kept under isoflurane anesthesia until the end of the surgery. To keep eyes moist an ocular lubricant was used. The scalp was shaved, disinfected with 70% alcohol and incised to expose the skull. Two small holes were drilled to insert a Nanofil needle (NF33BV) into the CeA (anterior/posterior, − 12 mm; medial/lateral, ± 26 mm; dorsal/ventral, − 50 mm, as measured from bregma and dura, Paxinos and Franklin 2008). Lentiviral vectors (4.5 × 10^7^ vector particles per site) or nanoparticles (0.5 μl total volume per site) were bilaterally injected into the CeA by pressure injection with a Nanofil syringe (10 μl) attached to the infusion pump (MicroSyringe Pump, World Precision Instruments; 100 nl/min). At the end of the injection the needle remained in place for additional 5 min for proper diffusion of the solution. All mice received tolfenamic acid (0.08%, 0.1 ml, 4.0mg/kg subcutaneous) and Enrofloxacin (Baytril 7.5 mg/kg) for 2 days after surgery (depending on their condition) and were left undisturbed for two days (nanoparticles) or one month (lentiviral vectors) before the start of behavioral experiments.

### Behavioral apparatus

All behavioral experiments were performed in the IntelliCage system, allowing for fully automated continuous testing of mouse behavior [15]. The apparatus consisted of a standard plastic cage (size 55 × 37.5 × 20.5 cm), with four operant learning chambers, each fitting into every corner of the housing cage, and a sleeping shelter in the center. The feeder was located above the shelter (food ad libitum). Each chamber was equipped with an infrared motion/proximity detector and a transponder reader that allowed identification of individual mice. Only one mouse could use the operant chamber at a time. Each chamber was equipped with two openings, each permitting access to a drinking bottle. Both openings were blocked with automatically operated motorized doors. Every event of a mouse poking its nose into the opening (a nosepoke response) and consumption of liquids (a lick response) were recorded for each mouse. Individual animals’ access to the bottles was assigned depending on the schedule of the experiment (described below). The system was controlled by a computer and monitored from the experimenter’s office via intranet. The mice were not disturbed, except for the technical breaks and exchange of bedding (once a week).

### Behavioral procedures

The mice were group-housed with a maximum of 10 animals per every IntelliCage. To identify individual mice in the IntelliCage, the animals were subcutaneously injected with sterile, glass-covered microtransponders (11.5 mm length, 2.2 mm diameter; Trovan, ID-100) under a brief isoflurane anesthesia. The experiment started with an adaptation to the new environment of the cage (simple adaptation, 4 days). In that condition the doors were open, and the animals had free access to water in all corners. Then, the doors in all corners were closed and access to water was gained after performing a nosepoke (nosepoke adaptation, 4 days). Next, mice had access to bottles limited to only one corner of the IntelliCage (place preference, 2 days). At the start of the discrimination training, the bottle less preferred during the place preference period was replaced with the bottle containing sweetened water (10% sucrose; appetitive discrimination training, 5 days). After the completion of the appetitive discrimination training mice had access to water in all four corners for 2 days. During the next 48 hours, mice had access to bottles only in one of the corners (which was different than the corner with the reward in the appetitive discrimination training). The bottle preferred during that phase was then replaced with a bottle containing quinine solution (0.005 M; bitter taste; aversive discrimination training, 5 days). Additionally, to assess the level of blocking of c-Fos expression, after the completion of the aversive discrimination training the mice had gained access to tap water in all four corners (2 days). Next, the animals were water deprived for 22 h to increase their motivation to seek the reward. Then, in the corner least preferred during the preceding phase the mice had access to two bottles that contained sweetened water (10% sucrose, on both sides of the corner). No access to water in other corners was offered. 90 minutes after completion of the training mice were sacrificed and their brains were taken for c-Fos expression analysis [8]. For c-Fos expression analysis after injection of nanoparticles releasing TIMP-1 and in MMP-9 KO mice the animals were exposed for 90 min to the novel cage containing a bottle with 10% sucrose solution and then sacrificed. Since we know that learning can be modulated by social context [18,19], the experimental and control groups were tested separately, with the similar numbers of animals per cage. The following numbers of animals were tested: LV_sh_c-fos (n=10), LV_sh_luc (n=9), TIMP-1 (two batches, n=7 and 7), CTRL (n=8), SYN-MMP9-GFP (n=10), SYN-GFP (n=10). After histological verification of injection sites the animals with misplaced injections were eliminated from further analysis. The final number of animals used in the analyses are given in the figure legends. To *assess colocalization of c-Fos and SST-Cre* 10 female C57Bl6 SST-Cre mice were exposed to the training in the IntelliCage (control, n = 4, reward-motivated training, n = 6, in one cage). Initially, the mice had access to the tap water in all corners. For the next 3 days, the animals were water deprived for 22 h to increase their motivation to seek the reward. The access to water was limited to 2h per day, at the beginning of the active phase [8]. Then, the sweetened water was placed in the corner, which was the least preferred during the previous step (10% sucrose solution, on both sides of the corner). Control mice were treated in a similar way, but they had access to tap water only (they were assigned to the corner containing tap water). Mice were sacrificed 90 minutes later and their brains were taken for c-Fos immunostaining procedure. To estimate MMP-9 activity in the CeA, prior to the training procedure, SST-Cre mice were injected with AAV-EF1a-DIO-EYFP-WPRE-pA vectors (n=4) to CeA. The animals were water deprived for 22 h, then in the reward-motivated group, mice were exposed to 10% sucrose solution, while the control animals drank only tap water. Mice were sacrificed 15 min after sucrose solution exposure and their brains were taken for measuring MMP-9 activity using synthetic, small peptide biomarker which is a substrate for MMP2/9 (A580, MMP Substrate 1, AnaSpec).

For the *whole-cell patch clamp recording* experiment mice were housed in pairs in their home cages (control, n=14, reward, n=11). For the next 3 days, the animals were water deprived for 22 h. Then, in the appetitive group, mice were exposed to 10% sucrose solution, 2h per day. After the last drinking session sucrose was replaced by tap water. Control mice were treated in the same way, but they drank tap water. Mice were sacrificed 18h after last sucrose exposure and their brains were taken for whole-cell recording.

### Immunohistochemistry

Mice were first anesthetized with isoflurane and then killed with an overdose of sodium pentobarbital (100 mg/kg) and perfused with 0,01M ice-cold PBS (Merck) followed by 4% PFA (Merck or Sigma-Aldrich). The brains were collected and fixed overnight in 4% PFA in PBS at 4°C. Then they were either immersed in 30% sucrose at 4°C and the tissue was frozen (isopentane, POCh on dry ice) and stored at −80C until sectioning into 40 μm sections on a cryostat (immunoenzymatic staining) or the brains were stored in PBS with 0,02% sodium azide (Sigma-Aldrich) at 4°C until sectioning into 40 μm sections on vibratome (immunofluorescent staining). Coronal brain sections containing amygdaloid nuclei (1.46–1.70 mm posterior to bregma, Paxinos and Franklin 2008) were collected. The immunohistochemical staining was performed on free-floating sections.

### Immunofluorescent staining

The brain slices were first washed free times in PBS, next permeabilized for 20 minutes in PBST (PBS with 0.3% Triton X-100; BioShop) and blocked for 2 h in 3% BSA (Sigma-Aldrich) in PBST at RT. Then, the sections were incubated with primary Polyclonal Guinea pig antibody (anti c-Fos, 1:1000; Synaptic Systems; Cat. No. 226 00), overnight at 4°C. After this, washed three times in PBS and incubated in secondary polyclonal antibody conjugated with Alexa Fluor^®^ 488 (goat anti-Guinea pig, 1:500, Invitrogen) for 2h in RT. Last, the sections were washed three times in PBS, mounted on slides and coverslipped with Fluoromount-G with DAPI (Invitrogen). Slides were imaged on ZEISS Spinning DISC confocal microscope. Regions of interest were imaged with 20x objective, z-stacks from the surface to 30 um in depth with 2um step. Later, all images were analyzed according to fluorescence in 488 channel for the c-Fos and 555 channel for the somatostatin cells with tdTomato marker. Image analysis was performed with a Java-based image processing program (ImageJ).

### MMP-9 activity detection

Mice were anesthetized with isoflurane and decapitated. The brains were removed and sliced using a Leica VT1200s vibratome in 4°C artificial cerebrospinal fluid (ACSF). Coronal slices of 150 μm thickness containing CeA were collected. Then, the sections were incubated for 30 minutes in ACSF with A580 MMP Substrate 1 (1 μg/ml, Anaspec) at RT. A580 MMP Substrate 1 is a small synthetic peptide (QXL 570 - KPLA - Nva - Dap(5 - TAMRA) - AR - NH2) in which fluorescence emission is blocked by intramolecular quenching (from TAMRA to QXL). When the peptide is cleaved by MMP2/9, intramolecular FRET is disrupted, causing emission of fluorescence following excitation at 545 nm. After incubation with biosensor, the slices were washed in ACSF and fixed in 4% PFA. Last, the sections were washed three times in PBS, mounted on slides and coverslipped with Fluoromount-G with DAPI (Invitrogen). Regions of interest were imaged with 63x oil objective on ZEISS Spinning DISC confocal microscope.

### Brain slice preparation for in vitro electrophysiology

18h after completing the behavioral procedure mice were anesthetized with isoflurane and decapitated. Then the brains were removed and sliced using a Leica VT1200s vibratome in 4°C artificial cerebrospinal fluid (ACSF) of the following composition of (in mM): 119 NaCl, 2.5 KCl, 2 MgSO_4_, 2 CaCl_2_, 1.25 NaH_2_PO_4_, 26 NaHCO_3_, 10 D-glucose equilibrated with 95% O_2_/5% CO_2_ gas composition. Coronal slices of 250 μm thickness containing CeA were collected.

### Whole-cell recording in acute brain slices

After 1–2 h of recovery, a single slice was transferred from a holding chamber to a recording chamber where it was continuously perfused with ACSF at RT. The whole-cell patch-clamp recordings were obtained from SST+ interneurons located in the lateral and medial division of the central amygdala. SST+ cells were identified by the presence of the fluorescent marker (tdTomato) expressed in a CRE-dependent manner. Recordings were performed using borosilicate glass electrodes (resistance 5-8 MΩ) filled with a solution composed of (in mM): 125 potassium gluconate, 2 KCl, 10 HEPES, 0.5 EGTA, 4 MgATP and 0.3 NaGTP, at pH 7.2, 290 mOsm. Data were acquired with a MultiClamp 700B amplifier (Molecular Device), filtered at 3 kHz, then digitized at 20 kHz and collected with pCLAMP software v. 10.07 (Molecular Device). Input resistance, neuronal capacitance and access resistance were monitored automatically. Intrinsic excitability was measured in current-clamp mode as a frequency of spiking in response to the SSTa current injection of the increasing intensity (500-ms square pulse, every 10 pA). The membrane potential of the cell was kept at the membrane potential of −65mV.

### Data analyses and Statistics

All results are presented as mean ± S.E.M. and were analyzed by the GraphPad 6 or 8.0.2. (Prism) and Statistica 7.0 software. *P* < 0.05 was considered significant (\**P* < 0.05, \*\**P* < 0.01, \*\*\**P* < 0.001).

No statistical methods were used to predetermine sample sizes, but our sample sizes are similar to those reported in previous publications (*10*). Normality tests were performed before choosing the statistical test. Wanting was analyzed with three-way ANOVAs with repeated measures. Liking and discrimination learning were analyzed with two-way ANOVAs. c-Fos expression in the blocking experiment was analyzed with one-way ANOVA. Numbers of c-Fos positive cells and double positive c-Fos+/SST+ cells were analyzed with a nonparametric Mann-Whitney U test. Electrophysiological experiments were analyzed using pCLAMP software (Molecular Devices). Kolmogorov-Smirnov test was applied to analyze statistical differences in the curves of Intensity-Frequency relationships between 2 groups of mice. The appropriate tests were chosen, taking into account whether data had a normal distribution and equal variation.

Injection sites, viral expression and nanoparticles presence were confirmed for all animals. Mice showing incorrect injection sites were excluded from the data analysis. Additionally, one mouse in c-Fos blocking experiment was removed from the analysis due to the lack of nosepoking activity during the adaptation phase of the behavioral experiments and 2 mice were not included in aversively motivated discrimination learning analysis because the data were not collected due to technical problems. In the MMP-9 overexpression experiments one mouse was eliminated from the control group due to the lack of nosepoking and licking activity.

## RESULTS

### Blocking c-Fos expression in the CeA impairs reward motivation and consumption, but does not influence respective aversive responses

Mice with local blockade of c-Fos expression in the CeA, achieved with lentivector carrying shRNA directed against *c-fos* mRNA (LV_sh_c-fos), and control animals injected with lentivector carrying shRNA directed against *luciferase* mRNA (LV_sh_luc, [16], Figure S1A), were subjected to discrimination training in the IntelliCages [9]. The shRNA injection resulted in reduction of c-Fos expression induced by the reward-motivated training in the CeA by about 50%, as compared to the control animals (Figure S1B). During the training, in one of the four corners of the IntelliCage, animals had to discriminate between two bottles (placed on two sides of the corner), which delivered either sweetened or tap water (appetitive task) or, alternatively, bitter (quinine-adulterated) or tap water (aversive task, Figure 1A). Approach/ avoidance motivation was measured as a number of nosepokes performed to the bottles with sweetened/quinine-adulterated water and tap water (Figure 1B,E), and consumption was measured as a percentage of licks of the sucrose/quinine solution to all recorded licks (Figure 1C,F). To assess spatial discrimination learning we measured a percentage of visits beginning with a correct response (to the bottle with sweetened water in the appetitively motivated task) or an incorrect response (to the bottle with quinine-adulterated water in the aversively motivated task) in the subsequent days of the training (Figure 1D,G). Blocking c-Fos expression in the CeA impaired all measured aspects of reward-motivated behavior, namely approach motivation, reward consumption, and appetitively motivated discrimination learning, leaving aversively-motivated behavior (discrimination learning, avoiding motivation, and consumption of quinine-adulterated water) intact. Importantly, the overall number of instrumental responses produced by the animals did not differ between the LV_sh_c-fos and LV_sh_luc groups neither in the appetitively nor in the aversively motivated task (data not shown).

**Figure 1.**
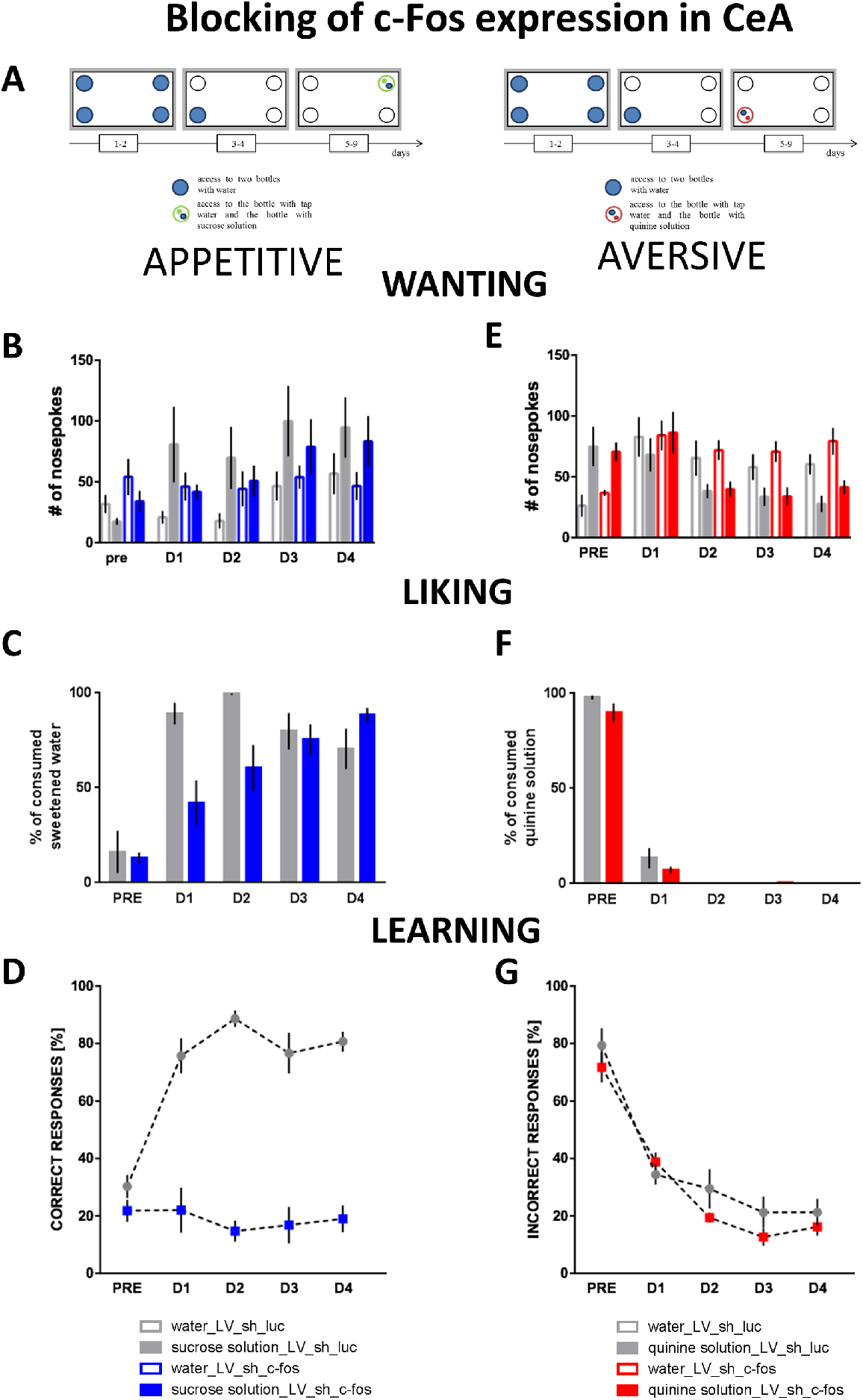
**A.** Schematic of the experiments on appetitive and aversive training in the IntelliCage. Prior to the training mice were injected with LV_sh_c-fos (n=6) or LV_sh_luc (n=6) vectors to CeA. **B.** Reward seeking (wanting), measured as a number of nosepokes made to obtain sweetened water, was diminished in LV_sh_c-fos mice [group x day x solution: F(4, 40)=3.15, p=0.0244]; # of nosepokes to water and sucrose differs at (at least) p<0.01 in D1-D3 in LV_sh_c-fos mice but not in LV_sh_c-fos mice (Bonferroni’s test). **C.** LV_sh_c-fos mice consumed less sucrose solution than LV_sh_luc mice during the first two days of the training [group: F (4, 40)=6.20, p=0.0006]; D1, D2: p<0.05 (Bonferroni’s test). **D.** Appetitively motivated discrimination learning was severely impaired in LV_sh_c-fos mice [group: F(1, 10)=96.60, p < 0.0001]. Avoidance motivation (**E**), consumption of quinine-adulterated water (**F**), and aversively motivated learning (**G**) were unaltered in LV_sh_c-fos mice. **pre**-adaptation period, **D1-4**, subsequent days of the training, bars denote SEM.

### Blocking MMP-9 activity in the CeA impairs reward motivation but not reward consumption, avoidance motivation, or bitter water consumption

Next, we measured approach and avoidance motivation, as well as sweetened and quinine-adulterated water consumption and learning in mice with MMP-9 activity locally inhibited in the CeA by tissue inhibitor of metalloproteinases-1 (TIMP-1) released from the PLGA nanoparticles (Figure S2A,B) [9,20]. Decreased activity of MMP-9 resulted in diminished approach motivation and impaired reward driven discrimination learning, however, consumption of sweetened water remained intact. Analogously to blocking c-Fos expression, decrease in MMP-9 activity did not affect the aversion towards or consumption of quinine-adulterated water, nor did it affect aversively motivated discrimination learning (Figure 2A-F). Of note, similar motivational changes were observed in mice globally lacking MMP-9 activity, because of the constitutional gene knock-out (MMP-9 KO, Figure S3).

**Figure 2.**
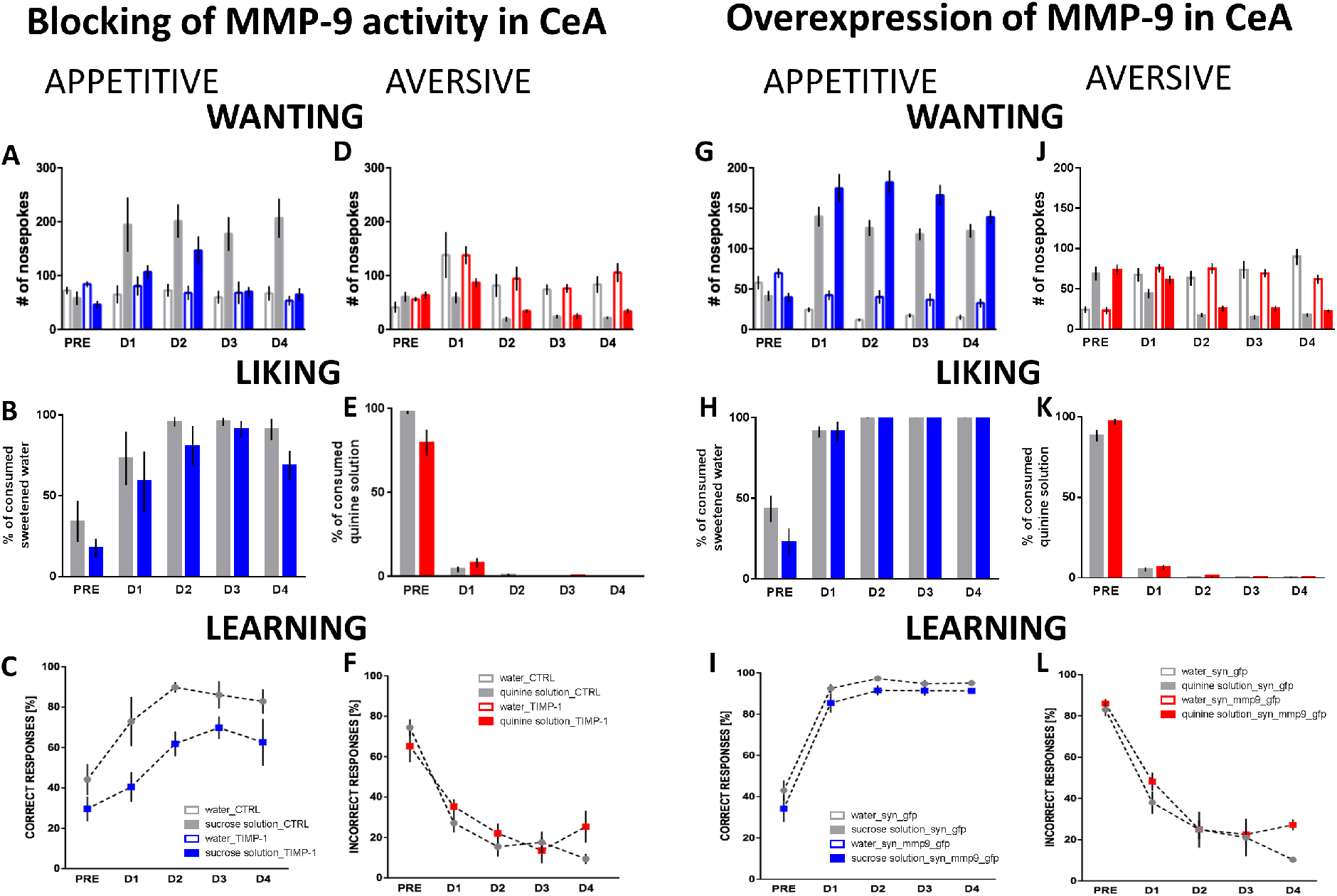
To inhibit MMP-9 activity in the CeA mice were injected with PLGA nanoparticles gradually releasing TIMP-1, an endogenous inhibitor of MMP-9 (n=6). Control animals (CTRL) were injected with nanoparticles carrying an inactive protein (BSA) (n=6). **A.** Reward seeking was diminished in TIMP-1-treated mice [group: F(1, 10)=12.62, p=0.0052]; # of nosepokes to water and sucrose differs at (at least) p<0.01 in D1-D4 only in CTRL mice (Bonferroni’s test). **B.** There was no difference in consumption of sucrose solution between the TIMP-1-treated and CTRL groups. **C.** Appetitively motivated discrimination learning was impaired in TIMP-1 mice [group: F (1, 10)=25.37, p=0.0005]. Avoidance motivation (**D**), consumption of quinine-adulterated water (**E**), and aversively motivated learning (**F**) were unaltered in TIMP-1-treated mice. To increase MMP-9 level in the CeA mice were injected with lentiviral vector carrying SYN-MMP9-GFP construct, causing MMP-9 overexpression (n=10). The control animals were injected with SYN-GFP construct (n=9). **G.** Reward seeking was increased when MMP-9 was overexpressed in the CeA [group: F(1, 17)=7.38, p=0.0147; group x day x solution: F(4, 68)=2.60, p=0.0435]. **H.** There was no difference in consumption of sucrose solution between the SYN-MMP9-GFP and SYN-GFP mice. **I.** Appetitively motivated discrimination learning was not changed in MMP-9-overexpressing group. Avoidance motivation (**J**), consumption of quinine-adulterated water (**K**), and aversively motivated learning (**L**) were unaltered in MMP-9 overexpressing mice. **pre**-adaptation period, **D1-4**, subsequent days of the training; bars denote SEM.

To confirm MMP-9’s role in the observed effects, we tested whether overexpression of MMP-9 in the CeA with lentiviral SYN-MMP9-GFP construct [17] also affects reward motivation. Injection of the construct resulted in a moderate overexpression of MMP-9 (Figure S2C). The MMP-9 overexpression did not significantly change consumption of sweetened water or reward-motivated discrimination learning. However, it clearly increased approach motivation. Notably, as found in case of blocking c-Fos expression and MMP-9 inhibition, increased MMP-9 levels affected neither aversion towards nor consumption of quinine-adulterated solution, nor aversively motivated discrimination learning (Figure 2G-L).

### Reward-motivated training induces c-Fos expression in both somatostatin-positive and somatostatin-negative neurons in the CeA but MMP-9 activation is associated with somatostatin-negative neurons only

To identify the subpopulation of CeA neurons expressing c-Fos as a result of the appetitively motivated training we initially checked colocalization of c-Fos with markers of inhibitory CeA neurons, VIP (vasoactive intestinal peptide), PV (parvalbumin) and SST (somatostatin). We analyzed c-Fos expression separately in the lateral and medial parts of the CeA, as they may differ functionally (7). We found that after the appetitively motivated training c-Fos is clearly expressed in the SST+ neurons, but only very sparsely in the VIP+ and PV+ neurons (Figure S4). The further experiments confirmed this result showing that in the lateral part of the CeA 55.5% ± 2.3% (mean ±SEM, n=6) of the c-Fos-expressing neurons were SST+, whereas in the medial part of the CeA 49.1% ± 1.4% (mean ±SEM, n=6) were c-Fos+SST+ (Figure 3).

**Figure 3.**
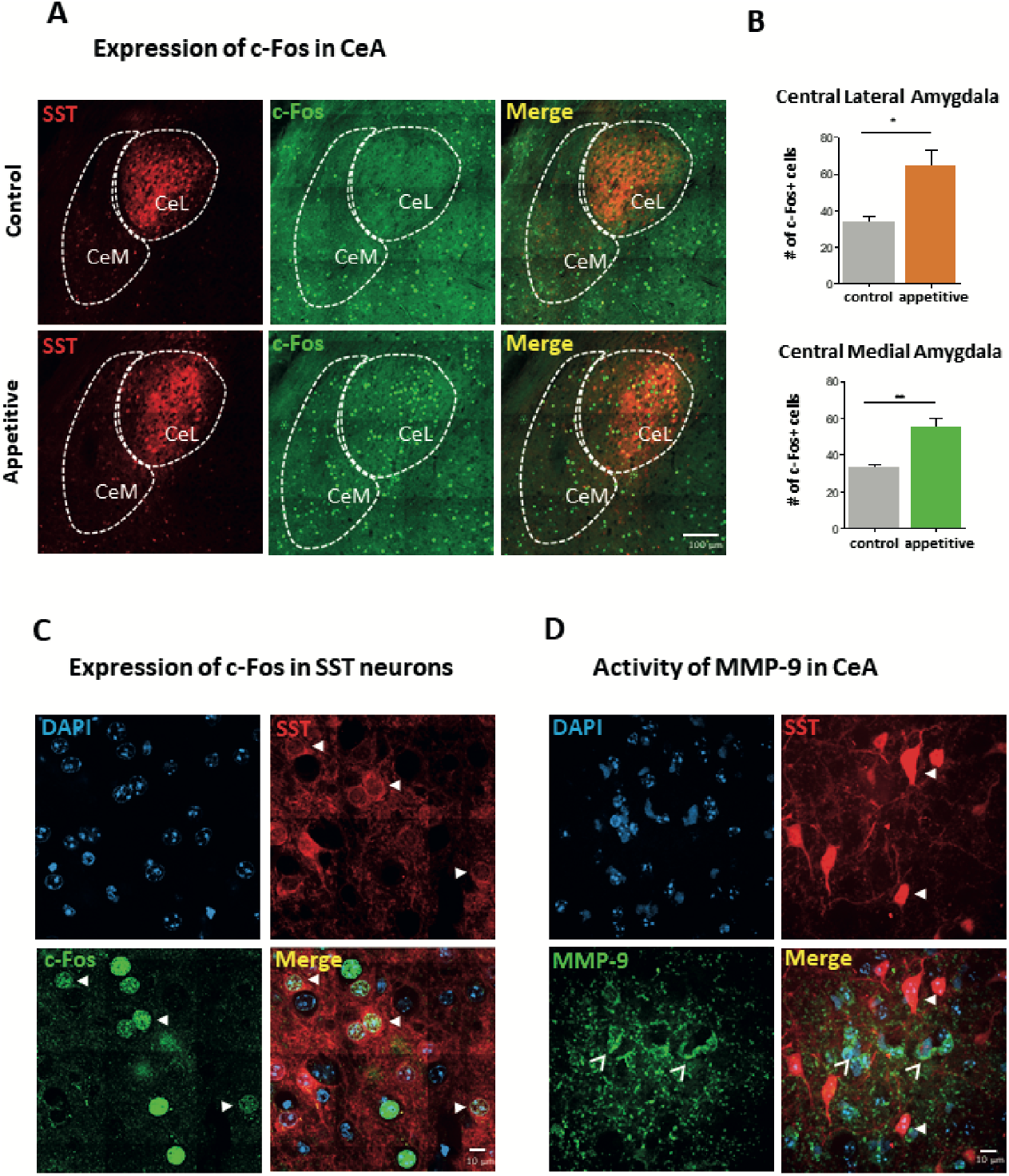
Expression of c-Fos and MMP-9 in the CeA induced by the reward-motivated training. **A.** Confocal images of coronal brain sections containing the CeA showing expression of c-Fos in SST+ and SST-cells in the SST-Ai14Tomato mice after reward-motivated training (Appetitive, n=6) and in control animals which drank only water in the IntelliCage (Control, n=4). Scale bar: 100 μm. **B.** The number of c-Fos-labeled cells in control animals and mice subjected to reward-motivated training in the medial (CeM) and lateral (CeL) parts of the CeA. Error bars represent SEM; significance level: \**p* < 0.05; \*\**p* < 0.01. **C.** Expression of c-Fos in SST+ neurons. Representative image showing CeA neurons, including SST+ cells expressing c-Fos induced by reward-motivated training. Scale bar: 10 μm. **D.** Activity of MMP-9 in the CeA. Confocal images of coronal brain sections containing CeA showing activation of MMP-9 in the CeA of SST-Cre mice injected with AAV-EF1a-DIO-EYFP-WPRE-pA for visualization of SST+ cells and then subjected to reward-motivated training. MMP-9 activation was detected using FRET biosensor A580 MMP Substrate 1. Please note that MMP-9 activity was detected only in SST-cells. Scale bar: 10 μm.

Next, to gain insight into the precise localization of training-driven MMP-9 activity we used A580 MMP Substrate 1, a small synthetic peptide in which fluorescence emission is blocked by intramolecular quenching. When the peptide is cleaved by MMP-9, intramolecular FRET (Förster Resonance Energy Transfer) is disrupted, causing emission of fluorescence following excitation at 545 nm. With this technique we observed increased MMP-9 activity in the CeA SST- cells of the SST-Cre mice subjected to appetitively motivated training in the IntelliCage (Figure 3). Thus, the results showed that after appetitively motivated training there were two subpopulations of cells, in which c-Fos is expressed, one comprised of the SST+ neurons that did not demonstrate MMP-9 activity, and another, consisting of the SST- neurons in which MMP-9 is activated.

### Reward consumption increases excitability of somatostatin-positive neurons in the CeA

The results of mapping of c-Fos and MMP-9 onto the SST neurons suggested that the c-Fos-MMP-9 pathway involved in reward wanting was associated with the SST- cells, while c-Fos-dependent liking is associated with the SST+ cells. To further test the latter, we studied electrophysiological properties of the SST+ cells after reward consumption. We modified the behavioral protocol so that it eliminated the reward seeking (wanting) component. For this reason the mice were exposed to the rewarding solution in their home cages and the drinking bottle, which usually contained tap water, was refilled with 10% sucrose solution. To test how reward consumption changes basic membrane features and the intrinsic excitability of the SST+ interneurons in the CeA, we performed whole-cell patch-clamp recordings in the lateral CeA (Figure 4).

**Figure 4.**
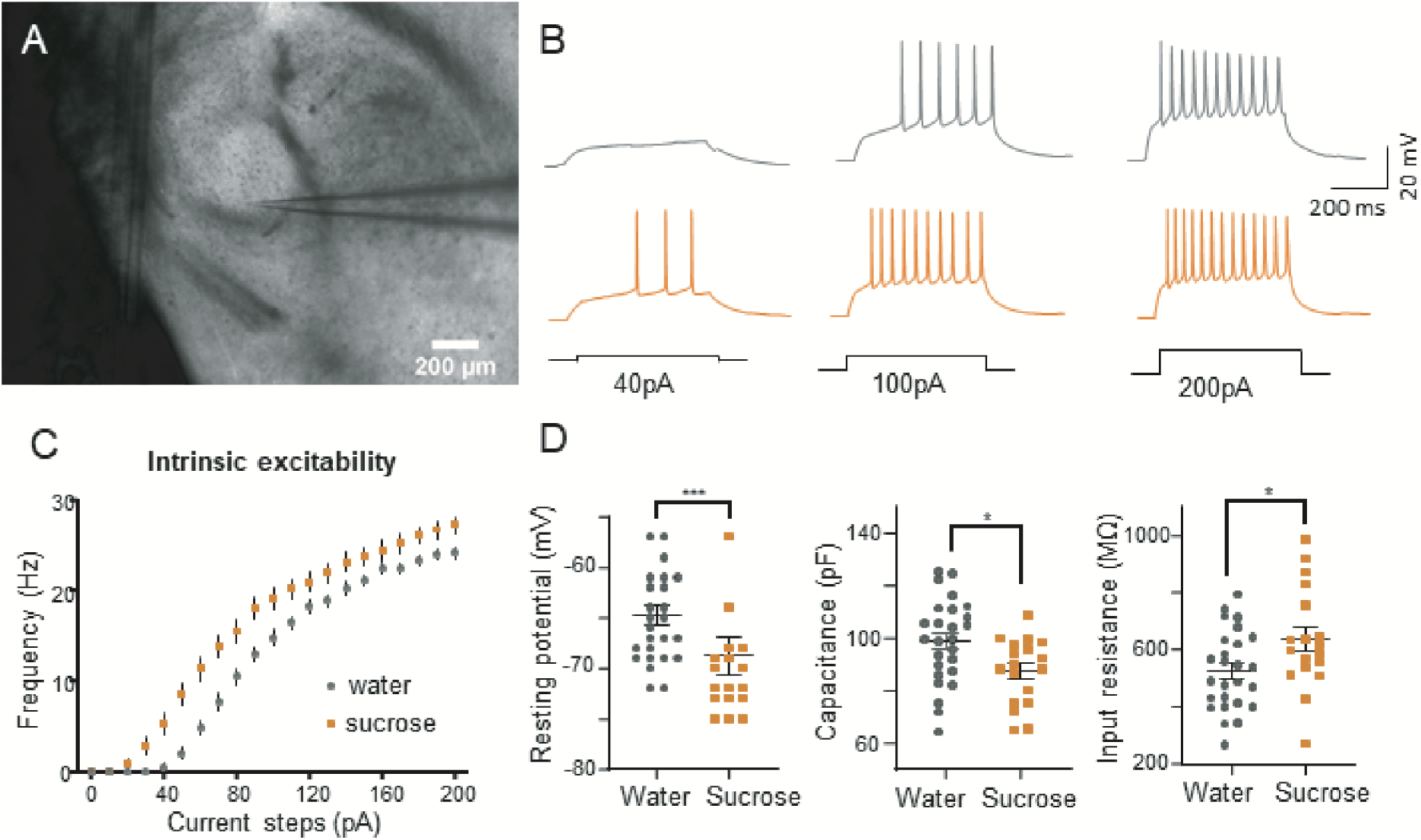
The effects of reward consumption on the electrophysiological properties of the SST+ neurons in the lateral subdivision of the CeA (CeL). **A**. Picture showing an acute brain slice with a whole-cell patch-clamp recording electrode located in the CeL. **B**. Majority of the SST+ neurons in the CeL are characterized by a delayed spiking (dLS) firing pattern (81%, tested in the control group drinking tap water). Representative traces of neuronal responses to three depolarizing current of increasing amperage are shown for the control (WATER, gray) and sucrose solution drinking (SUCROSE, orange) mice. **C.** Reward consumption lead to an increase in intrinsic excitability of the dLS SST+ neurons. Frequency of spikes in response to the increasingly depolarizing current steps injected to SST+ cells was higher in the SUCROSE (n=7) than in WATER (n=10) group (Kolmogorov-Smirnov test, p=0.0126). **D**. The effects of reward consumption on basic membrane properties of the dLS SST+ neurons: membrane resting potential (Mann-Whitney test, p=0.0010), membrane capacitance (Unpaired t test, p=0.0143) and input resistance (Unpaired t test, p=0.0254).

According to the shape of firing responses to somata current injection, 81% of the SST+ interneurons in control (water drinking) animals were characterized by delayed spiking (dLS, Figure 4B). We observed that reward consumption significantly lowered resting potential, as well as membrane capacitance, and raised membrane resistance in the dLS SST+ interneurons located in the lateral part of the CeA (Figure 4 D). We also found that reward consumption markedly increased intrinsic excitability in these interneurons (Figure 4B,C). In sum, the results show that reward consumption changes electrophysiological properties of the SST+ neurons.

## DISCUSSION

Here we report that motivation to pursue alimentary reward depends causally on c-Fos-MMP-9 pathway in the CeA. We show that blocking c-Fos-MMP-9 pathway disrupts approach motivation. At the same time inhibition of MMP-9 alone does not impair reward consumption. In contrast, reward consumption is modulated by c-Fos, which is expressed in somatostatin-positive subpopulation of the CeA neurons, in MMP-9-independent manner. Together, the results suggest that there are specialized molecular mechanisms associated with reward consumption, and pursuing alimentary rewards in the CeA, in line with the behavioral distinction between hedonic response to rewards and motivation to pursue them [3].

While discrimination learning is mediated by the cortico-striatal systems [21], the CeA has been implicated in modulation of incentive motivation to pursue a reward [22,23]. Our results further support that link. Molecular pathways related to appetitive motivation have not, however, been identified before. Recently, Kim et al. [24] described distinct groups of c-Fos positive CeA neurons responding to stimuli of positive vs. negative valence. The function of c-Fos expression and molecular mechanisms triggered by its activation in the CeA neurons have, however, been unclear. Our findings reveal the role of c-Fos-MMP-9 pathway in the modulation of approach motivation.

We show that blocking c-Fos expression disrupts both reward wanting and liking, while blocking MMP-9 activity impairs reward wanting, leaving liking intact. Thus, our results demonstrate the MMP-9-independent mechanism underlying reward consumption. Further, we identify the SST+ population of neurons in the CeA as relevant for those processes, by showing the increase in their excitability in response to reward consumption. It is noteworthy that the SST+ CeA neurons have been recently implicated in reward processing. For instance, activation of the SST+ lateral CeA neurons has been associated with abstinence-dependent expression of methamphetamine craving when drug-associated cues were presented during the relapse test [25]. The role of the CeA SST+ neurons in reward processing has been also suggested by the recent study showing that their inhibition results in depression-like behavior in a mouse model of chronic pain [26]. Since the CeA SST+ neurons send projections to the nucleus of the solitary tract and the parabrachial nucleus [27], the structures modulating taste-elicited responses and innervating the neurons projecting to the nucleus accumbens [28,29], the structure involved in goal-oriented behaviors, they are anatomically well positioned to modulate reward responses.

Our data suggest the existence of MMP-9-independent molecular pathway regulating consummatory aspects of reward behavior. Notably, whereas MMP-9 and TIMP-1 have emerged as major c-Fos/AP-1 gene targets in activated neurons, there is a number of other genes regulated by this transcriptional regulator that affect specific neuronal responses to external, behaviorally relevant stimuli. We argue that those other c-Fos/AP-1 targets may play a role in consummatory behavior (see, e.g., [30–33].

When blocking c-Fos expression, over days we observed a gradual improvement in instrumental behavior (nosepokes) aimed at obtaining reward, which suggests that encoding motivation was slowed down rather than blocked. This probably stems from the incomplete inhibition of c-Fos expression, since we reduced it to about 50% of the control level. On the other hand, inhibition of MMP-9 activity resulted in reduced instrumental behavior throughout the whole training, which is consistent with our previous reports [9] and *in vitro* tests of the nanoparticle-carried TIMP-1 activity [20]. Since TIMP-1 inhibits activity of not only MMP-9, but also other matrix metalloproteinases [30], to assess a specific role of MMP-9 in this phenomenon, we performed experiments in mice with a lentiviral vector-induced local overexpression of MMP-9 in the CeA [17]. As high levels of MMP-9 can disturb neuronal plasticity [34], we aimed at low to moderate overexpression (Fig. S2C), which indeed increased approach motivation leaving reward consumption intact and thus supported a bidirectional role of MMP-9 in motivational effects. One shall, however, consider that exogenously delivered MMP-9 might display activity outside of its physiological range.

To effectively elucidate a role of c-Fos and MMP-9 in the CeA in different subcomponents of reward-driven and aversively motivated behaviors we used the IntelliCage system, which offers a possibility of distinguishing between behaviors reflecting motivation, consumption and learning, similarly to the classical operant conditioning tests (9). The number of animals living in the cage was adjusted to exclude competition for the reward. Importantly, the animals were not food- or water-deprived during the tests, which could have affected the goal-directed behavior and c-Fos-dependent pathways [35,36]. The additional advantage of the IntelliCage is a possibility of long-term observation of voluntary behavior, in animals behaving at their own pace. Thus, such tests are particularly well-suited to study motivation, which is understood as willingness to take action.

Impaired motivation is a core feature of major depressive disorder, bipolar disorder, schizophrenia, as well as addiction [37–39]. In particular, habitual drug seeking has been shown to depend on the CeA [40]. Importantly, proteases have been shown to mediate experience-dependent plasticity both in animals [41] and in humans [42,43]. Since MMP-9 activity may be influenced pharmacologically [30,44], our results point to a potential target of intervention for motivational anhedonia.

## Authors contributions

Concept and design E.K., LK; data acquisition T.L., K.N., D.K., M.C., T.J., T.N., T.G., K.M.: analysis and interpretation of data T.L., K.N., J.D., D.K., J.J-S., T.N., K.M., J.U-C., E.K.; drafting and revising the article E.K., L.K., K.M.

## Data and materials availability

The datasets generated during and/or analyzed during the current study are available from the corresponding author.

## Supplementary Materials

### Supplementary methods

#### Animals

All procedures were carried out in accordance with the European Union and Polish guidelines for the care and use of laboratory animals, and were approved by the Local Ethical Committee. Female 3-4 month-old homozygous *mmp-9* knock-out mice on a C57BL/6J background (MMP-9 KO) and their wild-type (MMP-9 WT) littermates were used. The strain colony was maintained in the Animal House of the Nencki Institute. The animals were housed in a 12/12 h dark/light cycle, partially reversed (the dark phase shifted from 20:00–8:00 to 13:00–01:00 or 12:00–24:00 dependent on daylight savings), with food available ad libitum.

#### Immunoenzymatic staining

The sections were first washed three times in PBS, incubated for 10 min in 0.003% H_2_O_2_ in PBS, washed twice in PBS, and incubated with a polyclonal antibody (anti-c-Fos, 1:1000; Milipore, no. ABE457) and normal goat serum (3%; Vector) in PBS for 48 h at 4°C. Then, the sections were washed three times in PBST (PBS with 0.3% Triton X-100; Merck), incubated with goat anti-rabbit biotinylated secondary antibody (1:1000; Vector) and normal goat serum (3%) in PBST for 2 h at RT. The sections were then washed three times in PBST, incubated with avidin–biotin complex (1:1000 in PBST; Vector ABC kit) for 1 h at RT. The immunostaining reaction was developed using the oxidase-diaminobenzidine-nickel method. The sections were washed three times in PBS and incubated in distilled water with diaminobenzidine (DAB; Merck), 0.5 M nickel chloride for 5 min. The staining reaction was stopped by three washes with PBS. The reaction resulted in a brown-black stain within the c-Fos-positive nuclei of neurons. The sections were then mounted on slides, air dried, dehydrated in xylene, coverslipped with Entellan (Merck), and images containing CeA were collected with an optical microscope (Nikon Instruments). The c-Fos immunopositivity was measured as density, determined in the following manner. For each brain section, the number of c-Fos immunopositive nuclei in a given amygdalar structure was counted and divided by the area occupied by this structure (in mm^2^). Image analysis was done with the aid of an image analysis computer program (ImageJ) on at least two sections per animal.

#### Histology

Mice were killed by an overdose of sodium pentobarbital (100 mg/kg), perfused with ice-cold PBS (Merck) followed by 4% PFA (Merck). The brains were collected, fixed overnight in 4% PFA at 4°C, and subsequently immersed in 30% sucrose at 4°C. Then the brains were frozen and 40 μm sections were cut on a cryostat. The free-floating brain sections were collected in PBS. For injection site verification, the sections were directly mounted onto glass slides with VECTASHIELD mounting medium (Vector Laboratories) with 4,6-diamidino-2-phenylindole. A fluorescence microscope (Nikon Instruments) was used for imaging the brain sections.

#### In situ zymography

In situ zymography was done according to the protocol published earlier (Gawlak et al. 2009). In brief, coronal brain sections (4mm) containing the amygdalar complex were dissected immediately after the animals have been sacrificed and placed in methanol:ethanol mix (3:7, at 4°C) for 24 h. Then they were transferred to: 90% ethanol (1h at 4°C), 100% ethanol (2 x 1h, 4°C), 100% ethanol (30 min, 37°C), polyester wax (VWR 360704E):ethanol (1:3, 3h, 37°C), polyester wax:ethanol (1:1, ON+8h, 37°C), polyester wax:ethanol (9:1, ON, 37°C) and 100% polyester wax (ON, 42°C). Thereafter the tissue was placed at the bottom of a cylinder (ø 1cm) and embedded in fresh 100% polyester wax. When a wax column set (30 min in 4°C) it was placed, tissue facing upwards in a molten paraffin (Sigma Aldrich 327204) block precooled to about 42-45°C. When the paraffin block set (30 min in 4°C) it was attached to a wooden frame and subsequently was stored at −20°C. Sectioning into 6 μm coronal slices was done using a microtome equipped with the running water path (RMC, Boeckeler, Germany). Slides (Thermo Scientific Superfrost UltraPlus) with mounted brain slices were stored at 4°C until zymography in situ took place.

The in situ zymography reaction required de-waxing of the tissue (2×5 min +10 min in 100% ethanol at RT), followed by marking around with PAP Pen (Abcam ab2601) and irrigation with dH2O (3 x fast changes + 3×3 min). Preincubation in dH2O (100 min) took place in a wet chamber (37°C). The reaction with DQ gelatine (diluted 1:100 in the buffer supplied by the manufacturer, Invitrogen/Molecular Probes EnzChek™ Gelatinase/Collagenase Assay Kit E12055, 45 min) was also done in a wet chamber (37°C). It was followed by a wash-out in PBS (3×5 min, RT, Sigma, pH=7.4). Upon drying the slices were sealed with Fluoromount G (Invitrogen 00-4958-02) and left ON to dry before imaging on a Nikon (Ni-U) microscope.

---

**Figure S1.**
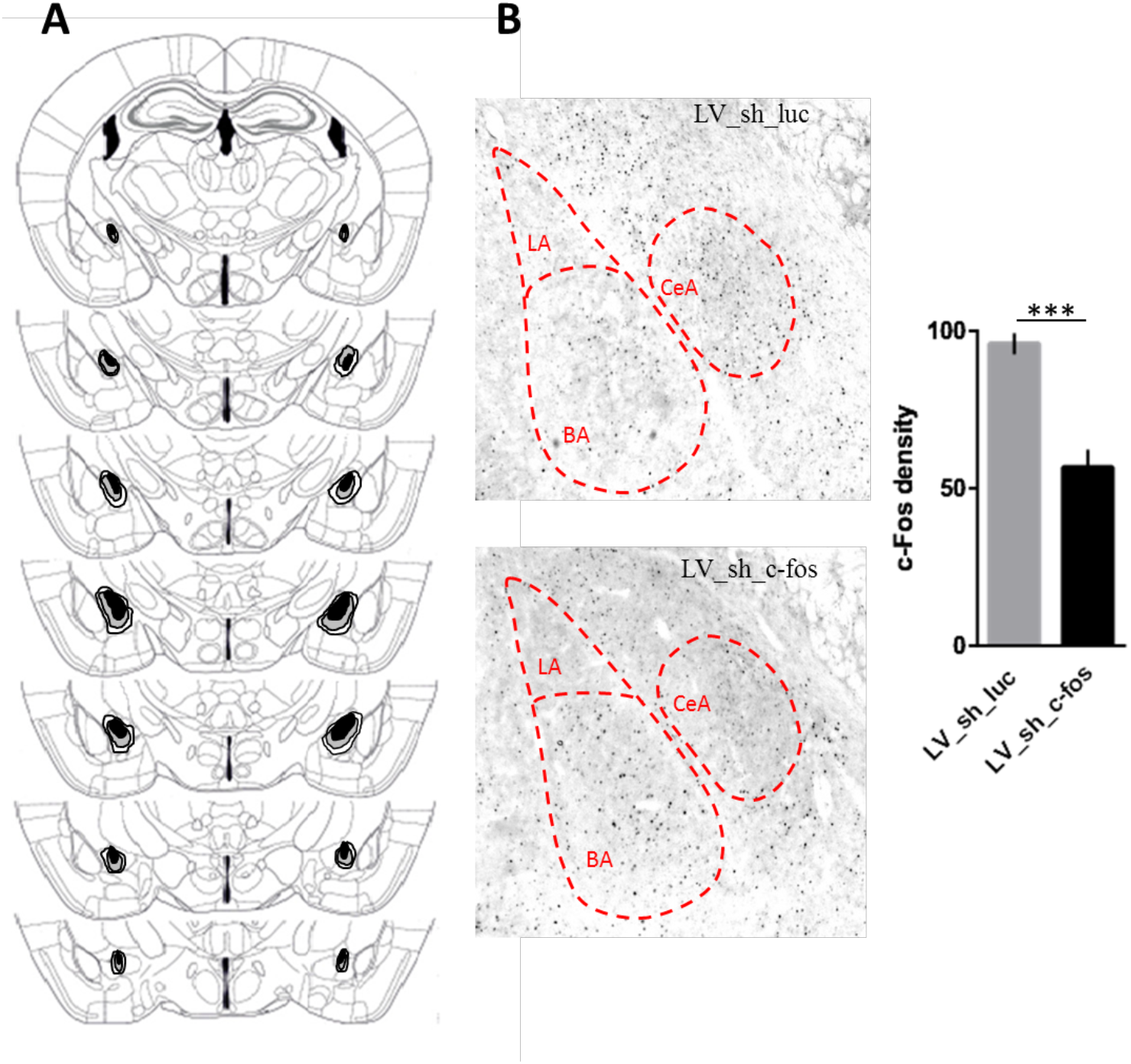
**A.** Prior to the training mice were injected with LV_sh_c-fos (n=6) or LV_sh_luc (n=6) vectors to CeA. Maximal (white), minimal (black) and average (gray) transfection range within the injection sites for animals included in the analysis are shown. **B.** Efficacy of *c-fos* blocking. Immunohistochemical staining for c-Fos in amygdala and quantified density of c-Fos expression in the region transduced by LV-sh-luc and LV-sh-fos vectors (F(1,10)= 35.78, p=0.0001). CeA - central, LA - lateral, BA - basal nuclei of amygdala; bars denote SEM, *** p<0.001.

**Figure S2.**
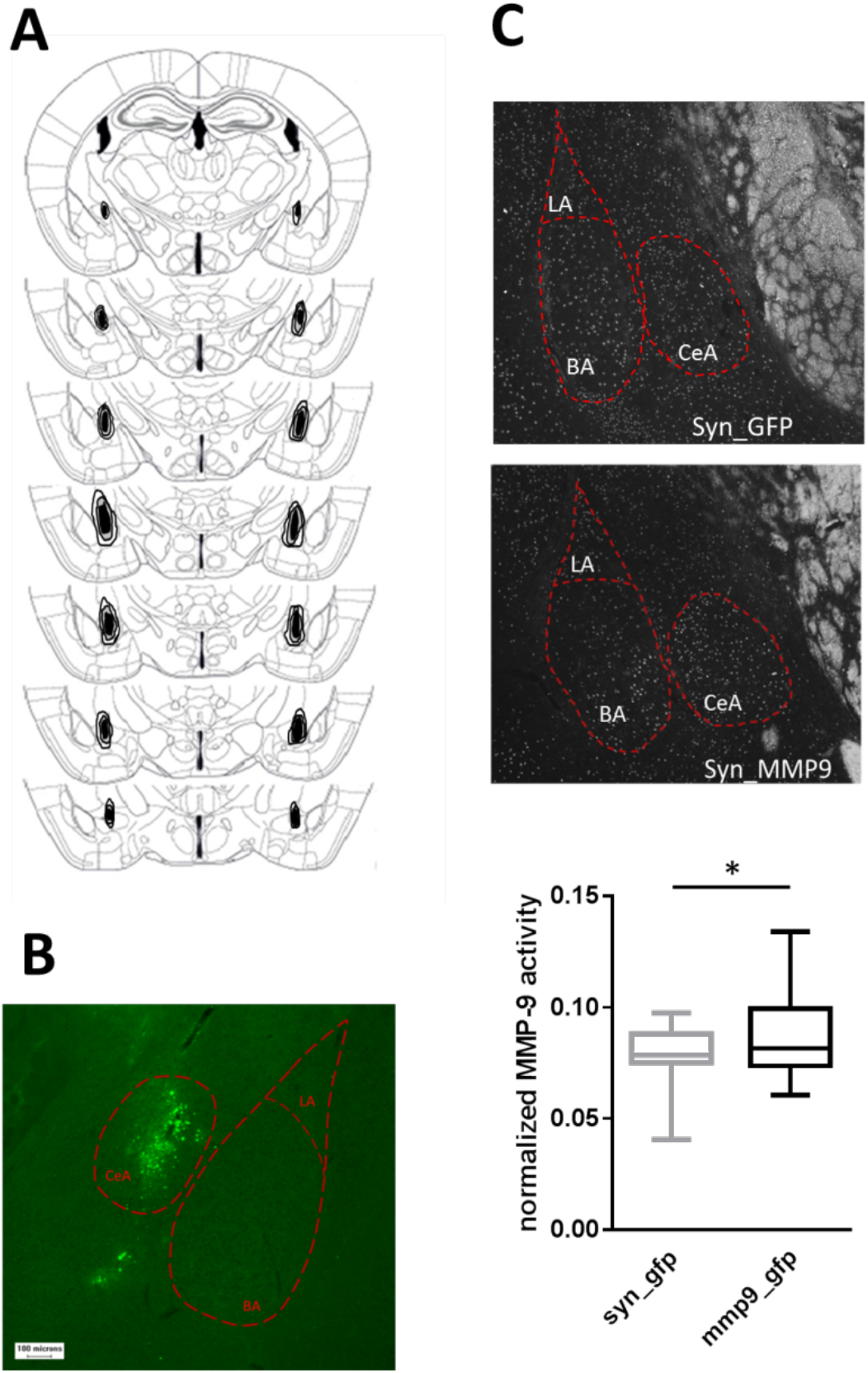
**A.** To inhibit MMP-9 activity in the CeA mice were injected with PLGA nanoparticles gradually releasing TIMP-1, an endogenous inhibitor of MMP-9 (n=6). Control animals (CTRL) were injected with nanoparticles carrying an inactive protein (BSA) (n=6). The diffusion of nanoparticles within the CeA is shown in the brain slice graphics; the largest (white), the smallest (black) and the average-sized (gray) injection sites for animals included in the analysis. **B.** An example of an injection site. **C.** To increase MMP-9 level in the CeA mice were injected with lentiviral vector carrying SYN-MMP9-GFP construct (n=10). The control animals were injected with SYN-GFP construct (n=9). In situ zymography (upper panel) showed a moderate increase in MMP-9 expression in the CeA of SYN-MMP9-GFP mice (min to max graph, K-S test, p=0.0275, lower panel). CeA - central, LA - lateral, BA - basal nuclei of amygdala, * p<0.05.

**Figure S3.**
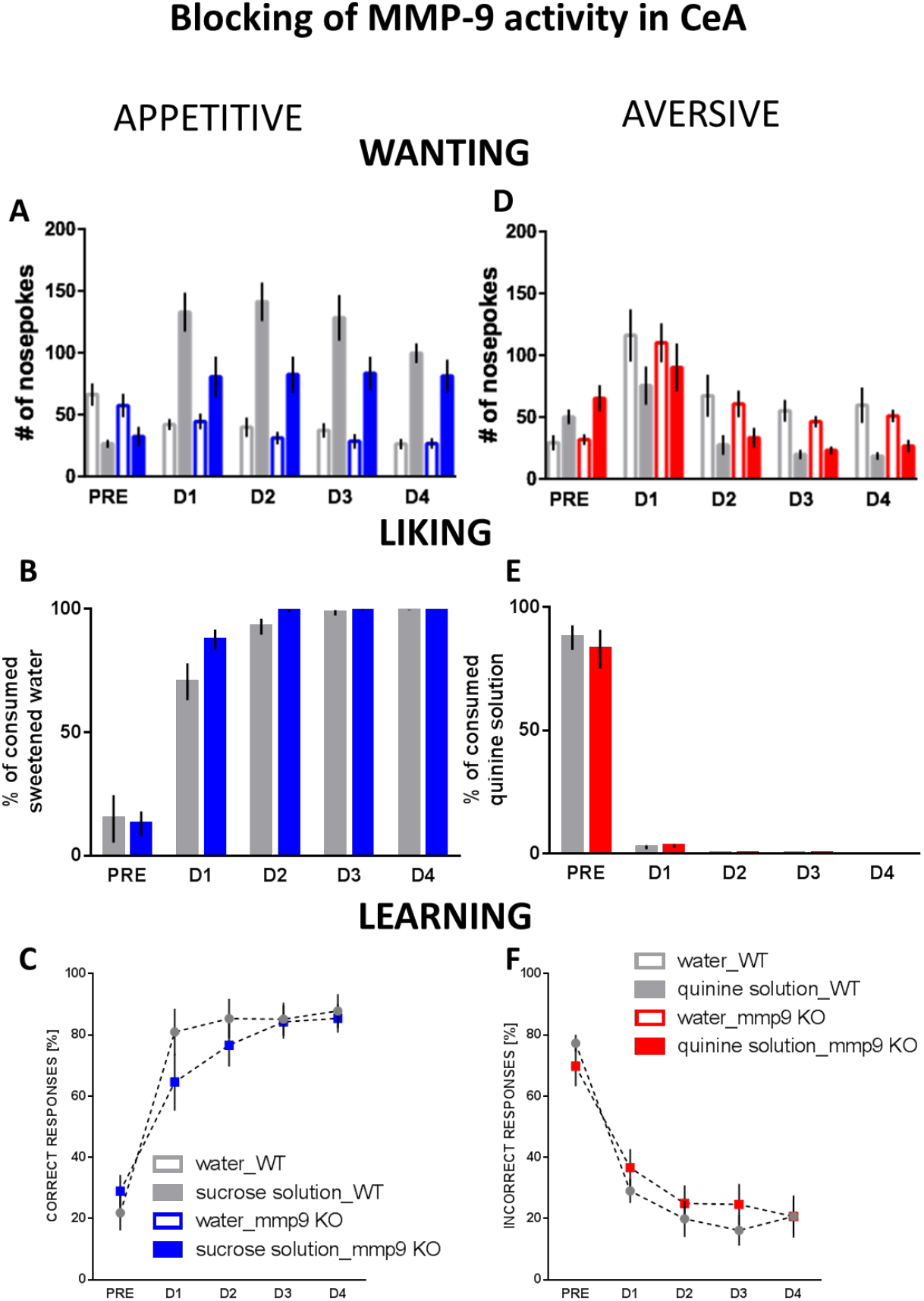
The effects of global lack of MMP-9 activity in the constitutive gene knock-out mice (MMP-9 KO n=12 and WT n=11). **A.** Reward seeking was diminished in the MMP-9 KO mice (group x session x solution interaction: F(4, 84)=4,28, p=0.0033). **B.** There was no difference in consumption of sucrose solution between the genotypes. **C.** Appetitively motivated discrimination learning was delayed in the MMP-9 KO group [group x session interaction: F (4, 84)=2.49, p=0.0490; distribution normalized with arcsin transformation]. Avoidance motivation (**D**), consumption of quinine-adulterated water (**E**), and aversively motivated learning (**F**) were unaltered in the MMP-9 KO mice. **pre**-adaptation period, **D1-4**, subsequent days of the training; bars denote SEM.

**Figure S4.**
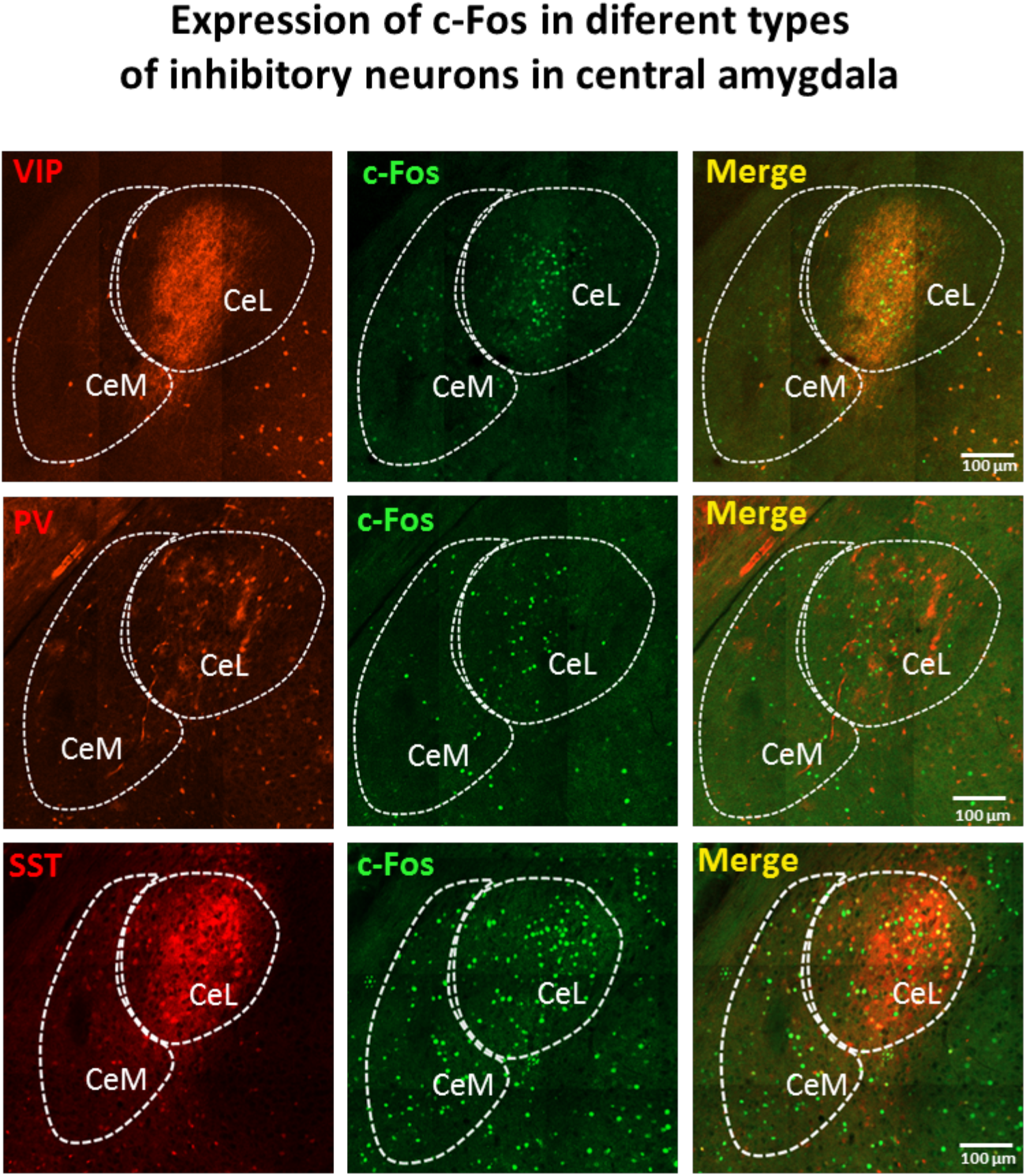
Representative images showing colocalization of c-Fos with markers of inhibitory CeA neurons, expressing vasoactive intestinal peptide (VIP), parvalbumin (PV) and SST somatostatin (SST) in animals subjected to appetitively motivated training in the IntelliCage. Please note that c-Fos is mostly expressed in the SST+ neurons, but only very sparsely in VIP+ and PV+ neurons. Scale bar: 100 μm.

